# Rhythmic variation in proteomics: challenges and opportunities for statistical power and biomarker identification

**DOI:** 10.1101/2024.06.28.601121

**Authors:** Matt Spick, Cheryl M Isherwood, Lee Gethings, Hana Hassanin, Daan R van der Veen, Debra J. Skene, Jonathan D Johnston

## Abstract

Time-of-day variation in the molecular profile of biofluids and tissues is a well-described phenomenon, but – especially for proteomics – is rarely considered in terms of the challenges this presents to reproducible biomarker identification. In this work we demonstrate these confounding issues using a small-scale proteomics analysis of male participants in a constant routine protocol following an 8-day laboratory study, in which sleep-wake, light-dark and meal timings were controlled. We provide a case study analysis of circadian and ultradian rhythmicity in proteins in the complement and coagulation cascades, as well as apolipoproteins, and demonstrate that rhythmicity increases the risk of Type II errors due to the reduction in statistical power from increased variance. For the proteins analysed herein we show that to maintain statistical power if chronobiological variation is not controlled for, *n* should be increased (by between 9% and 20%); failure to do so would increase β, the chance of Type II error, from a baseline value of 20% to between 22% and 28%. Conversely, controlling for rhythmic time-of-day variation in study design offers the opportunity to improve statistical power and reduce the chances of Type II errors. Indeed, control of time-of-day sampling is a more cost-effective strategy than increasing sample sizes. We recommend that best practice in proteomics study design should account for temporal variation as part of sampling strategy where possible. Where this is impractical, we recommend that additional variance from chronobiological effects be considered in power calculations, that time of sampling be reported as part of study metadata, and that researchers reference any previously identified rhythmicity in biomarkers and pathways of interest. These measures would mitigate against both false and missed discoveries, and improve reproducibility, especially in studies looking at biomarkers, pathways or conditions with a known chronobiological component.

## Introduction

Investigating time of day variation in different human biofluids and tissues has been a growing focus of research over the past decade. ^1^ This variation can be driven by endogenous circadian rhythms, which are influenced by central and/or peripheral clocks, as well as by exogenous diurnal rhythms resulting from behavioural and environmental cycles. In turn, dysregulation or disruption of these chronobiological variations has been associated to a wide range of diseases and conditions. ^2–6^ These rhythms are well described in terms of metabolomics and transcriptomics; ^7–10^ however, chronobiological variations in proteomic expression are less well mapped. ^11–13^ This is in part due to the sheer range of proteins and the relatively high costs of untargeted proteomic experiments; the precise number is unknown but a count of around 10,000 distinct proteins has been suggested for peripheral blood, ranging in concentration over at least 9 orders of magnitude. ^14^ This diversity has resulted in substantial gaps in current knowledge about which proteins and pathways exhibit diurnal or endogenous rhythmicity.

The rhythmicity of protein expression is relevant beyond analysis of circadian rhythm disorders, as temporal variation increases the potential for study design error and reduces statistical power. On the first issue, if a protein has a rhythmic component, this creates the potential for confounding if studies are designed without taking rhythmicity into account. This might occur, for example through selection / sampling bias, such as in a case / control study if all cases were to be measured in a hospital setting during a ‘morning round’, and controls were to be measured by convenience sampling, perhaps later in the working day. This bias would increase the risks of Type I errors, i.e. false positive identification of biomarkers as differentiating between cases and controls, with the risks of bias proportionate to the rhythm amplitude. Rhythmicity also increases the risk of Type II errors, by reducing statistical power. This reduction in statistical power for a given study follows naturally from the increase in variance in the features (here proteins) being measured. These challenges to study design can, however, also offer opportunities. By controlling for chronobiological variation through – for example – controlled time-of-day sampling, variance can be reduced and statistical power improved. Such study design steps are likely to be much more cost-effective than simply increasing the number of participants.

In this work, we provide a case study of the significance of rhythmicity for a small set of high-abundance proteins in human serum, by measuring protein concentration at a two-hourly resolution across 30 sequential hours and identifying rhythmic features alongside their acrophase (peak) values and amplitudes. We use these data to provide a real-world illustration of the challenges to study design that rhythmicity presents, in terms of increased *n* to maintain statistical power (or the opportunity to reduce *n* and maintain statistical power, if rhythmicity is controlled for). Finally, we set out four suggested mitigation steps against both Type I and Type II errors driven by chronobiology in proteomic analyses. These include incorporating time of sampling in study design; giving consideration to chronobiological issues in statistical power calculations; reporting time of sampling for cases and controls (with p-values) as metadata; and reporting any literature identified rhythmicity in any biomarkers of interest, to help provide context for readers.

## Method and Materials

### Experimental model and subject details

The samples analysed in this work were selected from a previous study as described in Isherwood *et al*. ^15^ Briefly, 24 healthy male volunteers were recruited by the Surrey Clinical Research Facility (CRF); the study was given a favourable ethical opinion from the University of Surrey ethics committee and all participants gave written informed consent. Participants’ blood was sampled during constant routine (60 blood draws: every 30 minutes for 30 sequential hours). Inclusion criteria included being male; aged between 18 and 35 years; BMI between 18 and 30 kg/m^2^ (inclusive) at screening; and habitual (at least 5 days per week) hours in bed per night between 7-9 hours which included a bedtime between 22:00-01:00 and waketime between 06:00-09:00. An ESS (Epworth Sleepiness Scale) score <9 was treated as indicative of normal range daytime sleepiness, an HÖ (Horne-Östberg; diurnal preference) score between 30-70 was taken as the normal range and a PSQI (Pittsburgh Sleep Quality Index) score <5 was considered indicative of satisfactory sleep quality and sleep. Shift workers involved in night work within the past six months and volunteers that had travelled across more than two time zones within a month before the study were excluded. Additional exclusion criteria were, smokers or nicotine products users within the 6 months prior to screening, and any regular use of medication known to influence circadian rhythm. A 10-day pre-laboratory routine, a previously validated standard for our human chronobiology experiments, was employed. ^16–18^ The full details of both the pre-laboratory protocol and the laboratory protocol are set out in Isherwood *et al*.,^15^ including allowed sleep windows, motion and light measurement via Actiwatches [Cambridge Neurotechnology, Cambridge, UK] and diet diaries, supplemented by continuous glucose monitors (CGMs) [Freestyle Libre 2, Abbott Laboratories Limited]. Compliance with the pre-laboratory routine was monitored by CGM confirmed compliance to the meal eating times. Upon admission in the CRF in the afternoon of Day 0, the participants were assessed to confirm compliance to the pre-laboratory routine and a review of medication, an alcohol breath test and a urine sample for analysis of cotinine and drugs of abuse were performed. Participants were supervised throughout the laboratory sessions by medical/clinical research staff. Laboratory environmental conditions and meals are again set out in Isherwood *et al*. ^15^

During the laboratory period 12 participants consumed hourly small meals throughout the waking period and 12 consumed two large daily meals. Following the diet regimes, all participants underwent a constant routine protocol with 30-60 min blood sampling. From the dataset of 12 individuals on the small-meal regime 11 were selected at random. The samples from the constant routine were taken forward for this proteomics analysis.

### Sample preparation

Human sera (50 µL) were aliquoted into fresh Eppendorf tubes prior to denaturing with 5 µL of 0.1% (w/v) RapiGest™ (Waters Corporation, Milford, MA) in 50 mM ammonium bicarbonate and incubated at 80°C for 45 min. Following incubation, 100 mM DTT (3 µL) was added and incubated for a further 30 mins at 60°C to reduce the proteins, before being alkylated with 200 mM iodoacetamide (3 µL) at room temperature for 30 min. Trypsin 1:50 (w/w) (Gold Mass Spectrometry grade, Promega, Madison, WI, USA) was added to each sample for proteolytic digestion and left incubating overnight at 37°C. TFA was added to a final concentration of 0.5% (v/v) to hydrolyse the RapiGest and heated for a further 45 min at 37°C, before centrifuging for 25 min at 18,000 g. The supernatant was collected and 3 µL aliquoted for LC-MS analysis. Aliquoted samples were diluted 1:250 (v/v) with 750 µL of 0.1% FA (v/v) and 19 µL of MassPREP™ Digestion Standard Mix 1 (Waters Corporation, Milford, MA) was added as an internal reference.

### LC-MS analyses

Peptides resulting from the tryptic digests were analyzed using the Evosep One EV-1000 (Odense, Denmark) coupled to a SYNAPT™ XS mass spectrometer (Waters Corp., Wilmslow, UK). Samples were loaded onto the Evosep tips as per the manufacturer’s instructions. Peptides were separated using the Evosep 60 SPD method, configured with a EV-1064 column. MS data were collected with the SYNAPT XS mass spectrometer, operated in positive electrospray ionisation (ESI) mode with a nominal resolution of 25,000 FWHM (V optics). The capillary voltage was 3.2 kV, cone voltage was 35 V and source temperature was set at 100 °C. Data were acquired over 50-2000 Da mass range with a scan time of 0.5 s. All mass spectral data were acquired in continuum mode using UDMSE to obtain fragmentation data simultaneously. ^19^ Function one (low energy) data were collected using a constant trap and transfer energy of 6 eV whilst the second (high energy) function consisted of a transfer collision energy ramp of 19 to 45 eV. For mass accuracy, [Glu1]-fibrinopeptide (*m/z* = 785.8426) was acquired as lock mass at a concentration of 100 fmol/µL (in 50:50 CH3CN/H2O, 0.1 % formic acid). Lock mass scans were collected every 60 s and averaged over 3 scans to perform mass correction. The time-of-flight was externally calibrated over the acquisition mass range (50-2000 Da) before analysis with a NaCI mixture. These data were collected using MassLynx v 4.1 software (Waters Corp., Wilmslow, UK) in a randomized order with three technical replicates acquired per sample. Lock mass consisting of [Glu1]-Fibrinopeptide was delivered to the reference sprayer of the MS source using the M-Class Auxillary Solvent Manager with a flow rate of 1 µL/min.

### Data analysis

The times of sampling were mapped to each participant’s dim light melatonin onset (DLMO) time by a previously described method. ^20^ Progenesis QI for Proteomics (Nonlinear Dynamics, Newcastle upon Tyne, UK) was used to process all LC-MS data. Retention time alignment, peak picking and normalization were conducted to produce peak intensities for retention time and *m/z* data pairs. Data were searched against reviewed entries of a *Homo sapiens* UniProt database (20,435 reviewed entries, release 2024) to provide protein identifications with a false discovery rate (FDR) of 1%. A decoy database was generated as previously described, ^21^ allowing for protein/peptide identification rates to be determined. Peptide and fragment ion tolerances were determined automatically, and searches allowed for one missed cleavage site. Carbamidomethyl of cysteines was applied as a fixed modification, whilst oxidation of methionines and deamidation of asparagine/glutamine were set as variable modifications. Following this process, a dataset in the form of an array comprising protein concentrations across features by time by participant was generated. These concentrations were then normalised by standard scaling, i.e. dividing by the per-participant standard deviation to express each concentration as a participant z-score. This generated an array of scaled protein concentrations with dimensions of participant *n* by DLMO time *t* by protein identifier *p*.

Cosinor analysis for rhythm detection was performed using CosinorPy (version 3.0) in the Python programming language (version 3.9.18), using the Spyder IDE (version 5.4.3). ^22,23^ A single component cosinor model was used in this work. CosinorPy includes functionality for multiple components, and also for inclusion of non-linear factors, but given the small sample size employed here, complex models were rejected due to the risk of overfitting. Proteins that were identified as rhythmic with p-value < 0.05 and amplitude > 0.1 were collated and processed for pathway upregulation / downregulation using the STRING online platform, ^24,25^ in order to identify clusters of proteins that were significantly altered (enrichment p-value < 0.05, with FDR correction).

To assess the impact of rhythmicity on biomarker identification and statistical power, we start with the assumption that the variance of a biomarker measured in a cohort controlled for chronobiological variation is 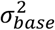. To this base variation, an uncontrolled chronobiological variation represented by a cosine function would add additional variance 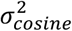, plus the covariance 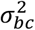, as shown in eq. 1. If there is no relationship between base variation and variation due to chronobiology, the covariance term is zero. Furthermore, the chronobiological variance could be expressed solely in terms of the amplitude *A* of the cosine function, yielding eq. 2. For biomarkers with multiple components, total variance will be the sum of the base variance (excluding any rhythmic component) plus the sum of *A*^*2*^/2 for each additional cosinor component (by the variance sum law). ^26^

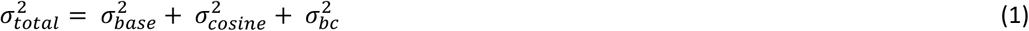

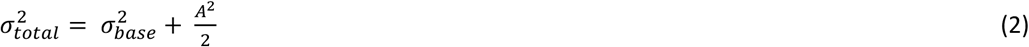

Because variance increases when adding uncorrelated cosinor components, the number of participants *n* required to maintain statistical power would be increased for given critical values *Z* and effect size *d*, as illustrated in eq. 3. The critical values used here were based on *α* = 0.05 and β = 0.20.

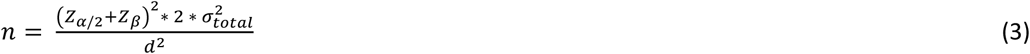

For each rhythmic protein identified here, the increase in *n* required to maintain statistical power is reported. Additionally, the impact on power (*Z*β and therefore β) is derived from eq. 4).

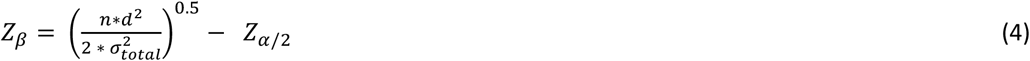

## Results

### Cohort analysis

11 participants were randomly selected from the small meal cohort of 12 from the original study; 1 was excluded from subsequent analysis due to failed LC-MS injections. The key characteristics of the 10 study participants analysed in this work are summarised in Table 1. All recruited participants were male.

**Table 1:** Characteristics of study participants (n = 10)

|  | Mean | Standard Deviation |
| --- | --- | --- |
| 25% DLMO (dec. H) | 22.77 | 1.11 |
| Age (years) | 28.2 | 4.3 |
| Height (cm) | 180 | 7.8 |
| Weight (kg) | 78.6 | 9.3 |
| BMI (kg m<sup>-2</sup>) | 24.3 | 2.6 |
| Body fat (%) | 14.7 | 5.5 |
| HO (score) | 58.4 | 4.7 |
| PSQI (score) | 3.5 | 0.5 |
| ESS (score) | 3.5 | 2.6 |

### Rhythm analysis

Proteins (n = 281) were identified across the triplicate injections. All proteins with greater than 10% missing values were excluded from further analysis, leaving 73 proteins for rhythm analysis. Of these, 13 proteins were identified as including a 24-hour or 12-hour rhythm based on two criteria: a p-value when fitted to a single component cosinor of < 0.05 and an amplitude > 0.1. Of the 13 proteins, 9 showed a 12-hour rhythm and 6 showed a 24-hour rhythm, with an overlap of 2 proteins. The individual proteins identified as rhythmic are shown in Table 2 with their acrophases and amplitudes. These amplitudes are shown in terms of the intra-individual z-scores, i.e. relative to the standard deviation exhibited by each individual.

**Table 2:**
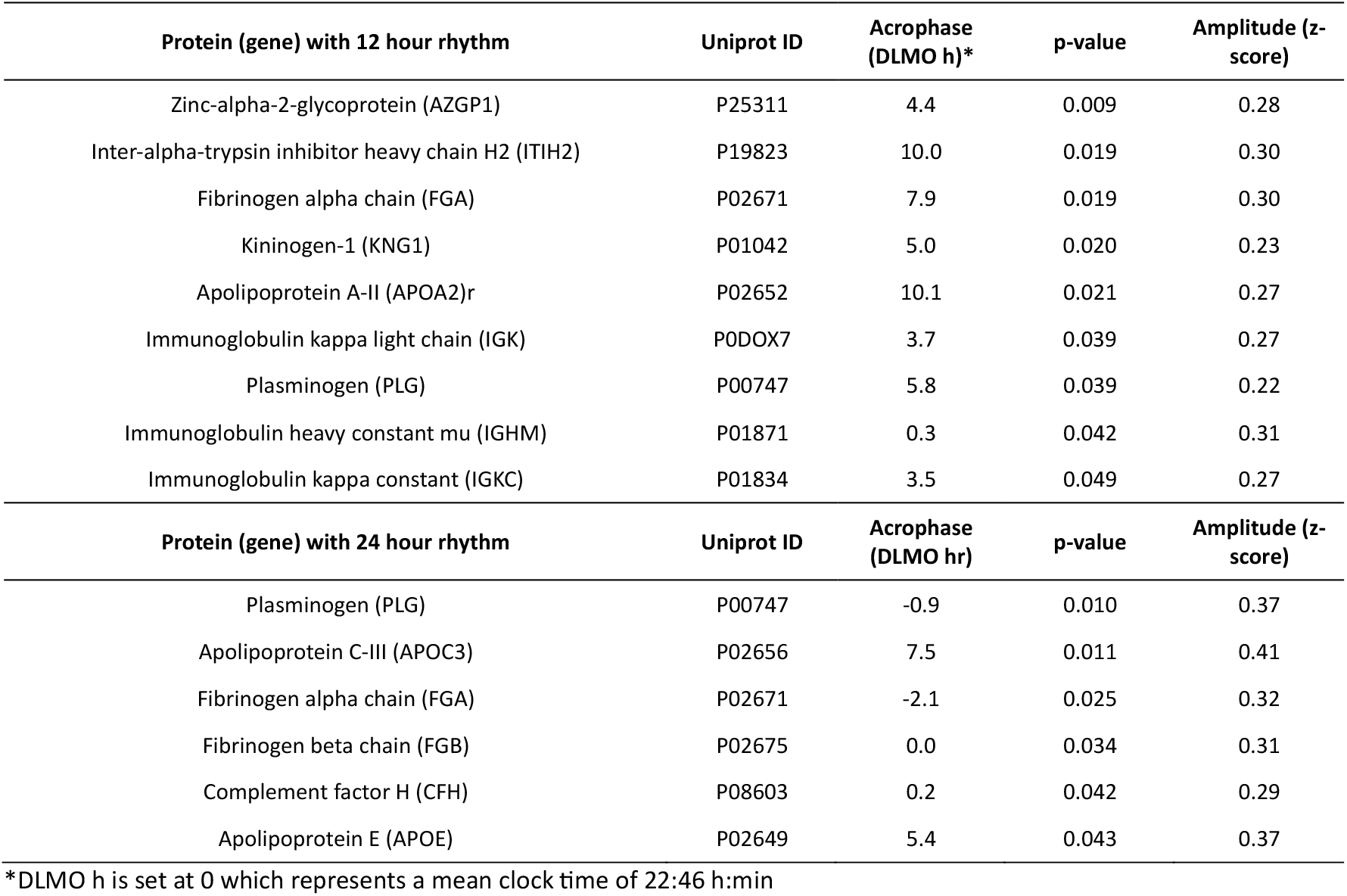
Cosinor analysis of proteins identified as having a significant rhythm (p < 0.05) versus H0 of no rhythmicity.

| Protein (gene) with 12 hour rhythm | Uniprot ID | Acrophase (DLMO h)* | p-value | Amplitude (z-score) |
| --- | --- | --- | --- | --- |
| Zinc-alpha-2-glycoprotein (AZGP1) | P25311 | 4.4 | 0.009 | 0.28 |
| Inter-alpha-trypsin inhibitor heavy chain H2 (ITIH2) | P19823 | 10.0 | 0.019 | 0.30 |
| Fibrinogen alpha chain (FGA) | P02671 | 7.9 | 0.019 | 0.30 |
| Kininogen-1 (KNG1) | P01042 | 5.0 | 0.020 | 0.23 |
| Apolipoprotein A-II (APOA2)r | P02652 | 10.1 | 0.021 | 0.27 |
| Immunoglobulin kappa light chain (IGK) | P0DOX7 | 3.7 | 0.039 | 0.27 |
| Plasminogen (PLG) | P00747 | 5.8 | 0.039 | 0.22 |
| Immunoglobulin heavy constant mu (IGHM) | P01871 | 0.3 | 0.042 | 0.31 |
| Immunoglobulin kappa constant (IGKC) | P01834 | 3.5 | 0.049 | 0.27 |
| Protein (gene) with 24 hour rhythm | Uniprot ID | Acrophase (DLMO hr) | p-value | Amplitude (z-score) |
| Plasminogen (PLG) | P00747 | -0.9 | 0.010 | 0.37 |
| Apolipoprotein C-III (APOC3) | P02656 | 7.5 | 0.011 | 0.41 |
| Fibrinogen alpha chain (FGA) | P02671 | -2.1 | 0.025 | 0.32 |
| Fibrinogen beta chain (FGB) | P02675 | 0.0 | 0.034 | 0.31 |
| Complement factor H (CFH) | P08603 | 0.2 | 0.042 | 0.29 |
| Apolipoprotein E (APOE) | P02649 | 5.4 | 0.043 | 0.37 |

\*DLMO h is set at 0 which represents a mean clock time of 22:46 h:min

Both the 24-hour and 12-hour sets of proteins were subjected to separate pathway analysis using STRING. For the 12-hour set of proteins, one cluster of proteins was identified (Figure 1A) with a p-value of < 0.001 for the overall level of enrichment. The proteins within this cluster were investigated for functional enrichment. Different databases may produce different results, but here showed considerable overlap; enrichment according to GO Biological Processes was significant for fibrinolysis (p=0.020), negative regulation of blood coagulation (p=0.004) and blood coagulation (p=0.020); whilst according to KEGG Pathways, functional enrichment was significant for the complement and coagulation cascades (p < 0.001). The full lists of pathways are shown in Table S1, Supplementary Material. Comparison of acrophases for the proteins identified also showed clustering for PLG, KNG1 and AZGP1 around 5 hours after DLMO (= 03.77 (dec. h, clock time) (Figure 1B).

**Figure 1:**
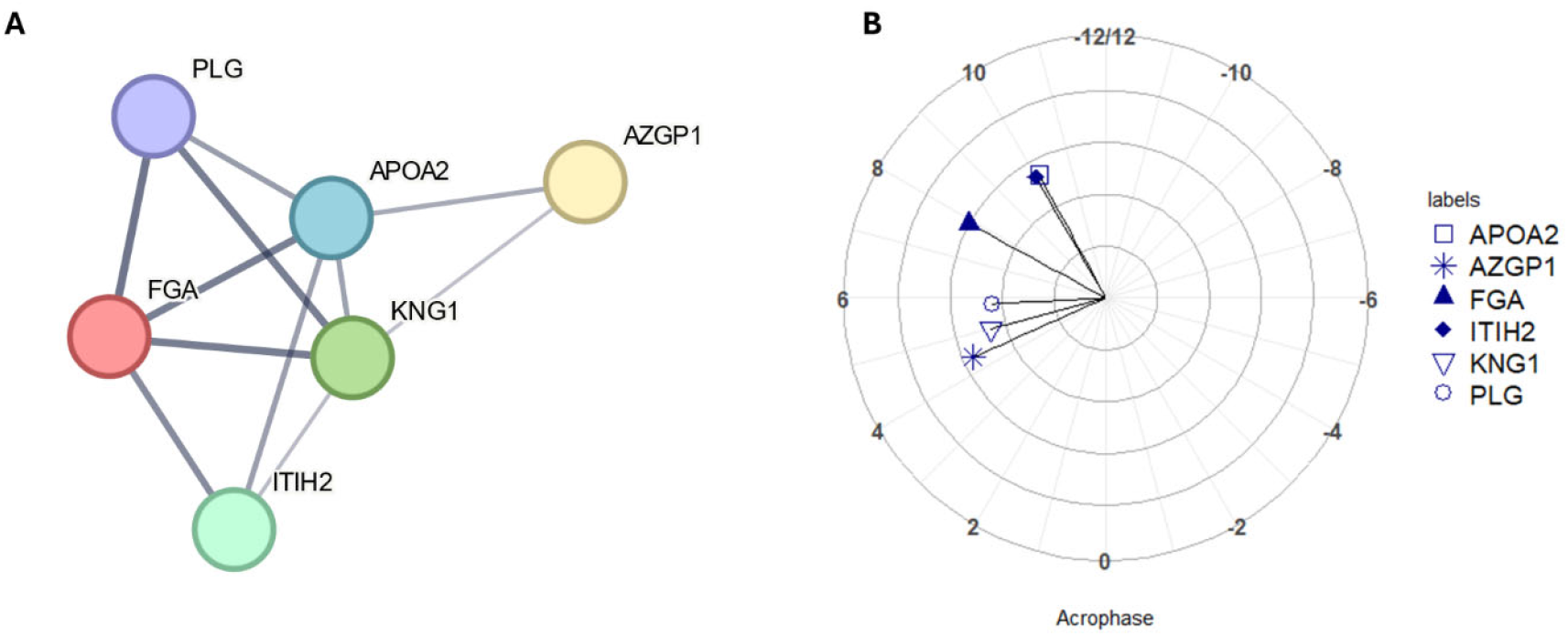
(A) Node and edge relationships of proteins identified as having statistically significant 12-hour rhythms (p-value < 0.05) and forming a functionally enriched cluster. Network nodes represent proteins and edges represent protein-protein interactions, line thickness indicates the strength of data support (B) polar plot of acrophases of proteins shown in Figure 2A, amplitude is shown in terms of z-score along the radius, DLMO time is shown around the perimeter (DLMO 0 = clock time 22:46 h:min). All analyses based on a cohort of n = 10.

For the 24-hour set of rhythmic proteins, one cluster of proteins was identified (Figure 2A) with a p-value of p < 0.0001 for the overall level of enrichment. The proteins within this cluster were investigated for functional enrichment in the same way as for the 12-hour set of proteins. Enrichment according to GO Biological Processes was significant for pathways including plasminogen activation (p=0.004), negative regulation of triglyceride metabolism (p=0.004); whilst for KEGG pathways, functional enrichment was again significant for the complement and coagulation cascades (p<0.001), for cholesterol metabolism (p=0.016) and platelet activation (p=0.049). The full lists of pathways are shown in Table S2, Supplementary Material. Comparison of peak timings showed that complement and fibrinolysis-related proteins all had an acrophase between a DLMO time of –2 hours and 0 hours (clock time 20.77-22.77 h)(Figure 2B).

**Figure 2:**
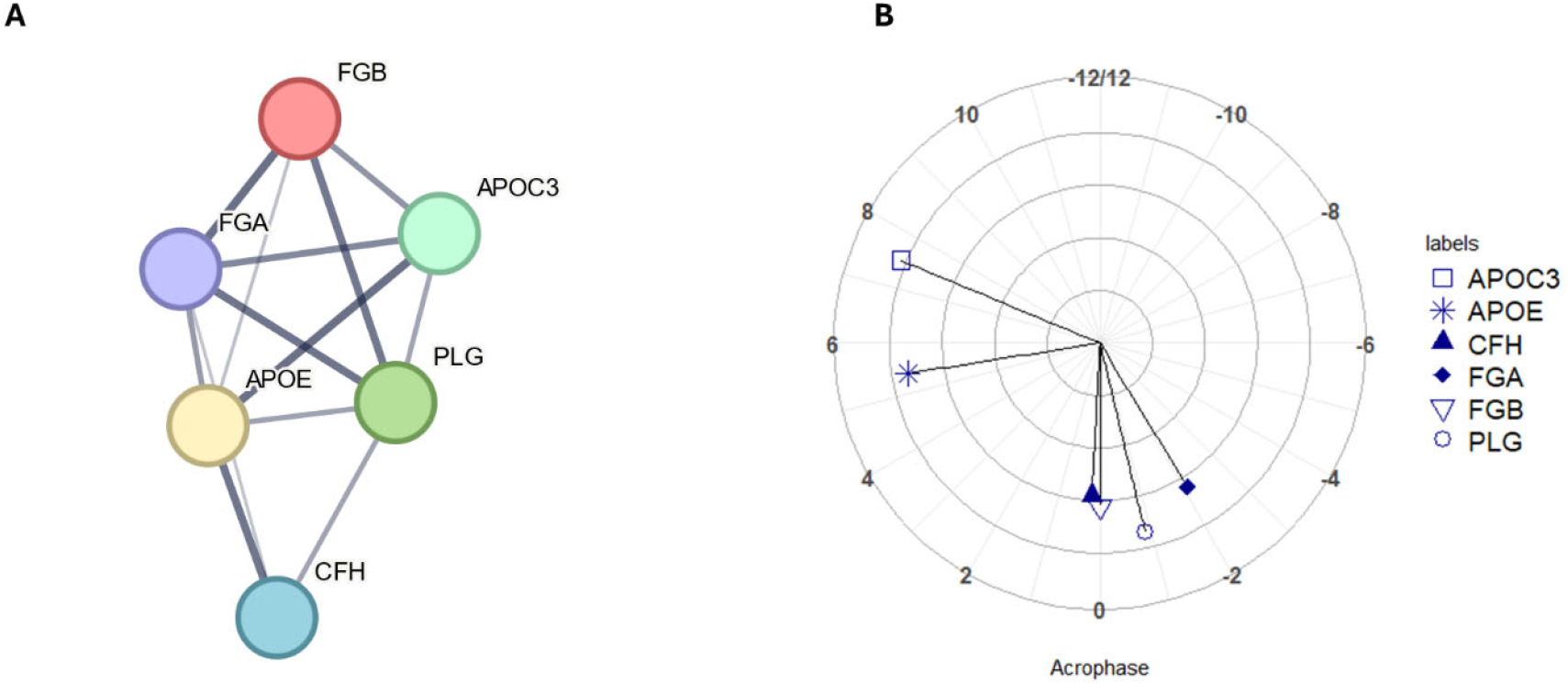
(A) Node and edge relationships of proteins identified as having statistically significant 24-hour rhythms (p-value < 0.05) and forming a functionally enriched cluster. Network nodes represent proteins and edges represent protein-protein interactions, line thickness indicates the strength of data support (B) polar plot of acrophases of proteins shown in Figure 3A, amplitude is shown in terms of z-score along the radius, DLMO time is shown around the perimeter (DLMO 0 = clock time 22:46 h:min). All analyses based on a cohort of n = 10.

Finally, the potential impact of rhythmicity on biomarker identification and pathway contribution is summarised in Table 3. This table shows for each protein the amplitude identified in this work, and the increase in *n* required to maintain statistical power if rhythm is not controlled, or conversely the decrease in *n* achievable for a given Type II error rate if rhythm is controlled.

**Table 3:** Statistical implications for proteins identified as rhythmic in this work.

| Protein (gene) with 12 hour rhythm | Amplitude (z-score) | Increase in *n* for maintained power | β (Type II error rate) if uncontrolled <sup>1</sup> |
| --- | --- | --- | --- |
| Zinc-alpha-2-glycoprotein (AZGP1) | 0.28 | 9% | 23% |
| Inter-alpha-trypsin inhibitor heavy chain H2 (ITIH2) | 0.3 | 10% | 24% |
| Fibrinogen alpha chain (FGA) | 0.3 | 10% | 24% |
| Kininogen-1 (KNG1) | 0.23 | 6% | 22% |
| Apolipoprotein A-II (APOA2) | 0.27 | 8% | 23% |
| Immunoglobulin kappa light chain (IGK) | 0.27 | 8% | 23% |
| Plasminogen (PLG) | 0.22 | 5% | 22% |
| Immunoglobulin heavy constant mu (IGHM) | 0.31 | 11% | 24% |
| Immunoglobulin kappa constant (IGKC) | 0.27 | 8% | 23% |
| Protein (gene) with 24 hour rhythm | Amplitude (z-score) | Increase in *n* for maintained power | β (Type II error rate) if uncontrolled |
| Plasminogen (PLG) | 0.37 | 16% | 26% |
| Apolipoprotein C-III (APOC3) | 0.41 | 20% | 28% |
| Fibrinogen alpha chain (FGA) | 0.32 | 11% | 24% |
| Fibrinogen beta chain (FGB) | 0.31 | 11% | 24% |
| Complement factor H (CFH) | 0.29 | 9% | 24% |
| Apolipoprotein E (APOE) | 0.37 | 16% | 26% |

<sup>1</sup> For baseline calculations a β of 20% was used, or a statistical power of 80%

### Power and Implications for Statistical Errors

In addition to the specific proteins analysed here, the data also provide a case study for calculating the impact of biological rhythmicity on statistical power. Both the Type II error rate and the number of participants required increase as the amplitude of the rhythmic component increases. The overall relationship is shown in Figure 3. For any given amplitude, there will be a required increase in *n* needed to maintain statistical power (Figure 3A), or an increased Type II error rate if *n* is not increased (Figure 3B). Conversely, controlling for rhythmicity allows for the maintenance of statistical power without increasing n. Both functions (for n and for the Type II error rate) vary with the square of the amplitude, so both charts show the same exponential relationship.

**Figure 3:**
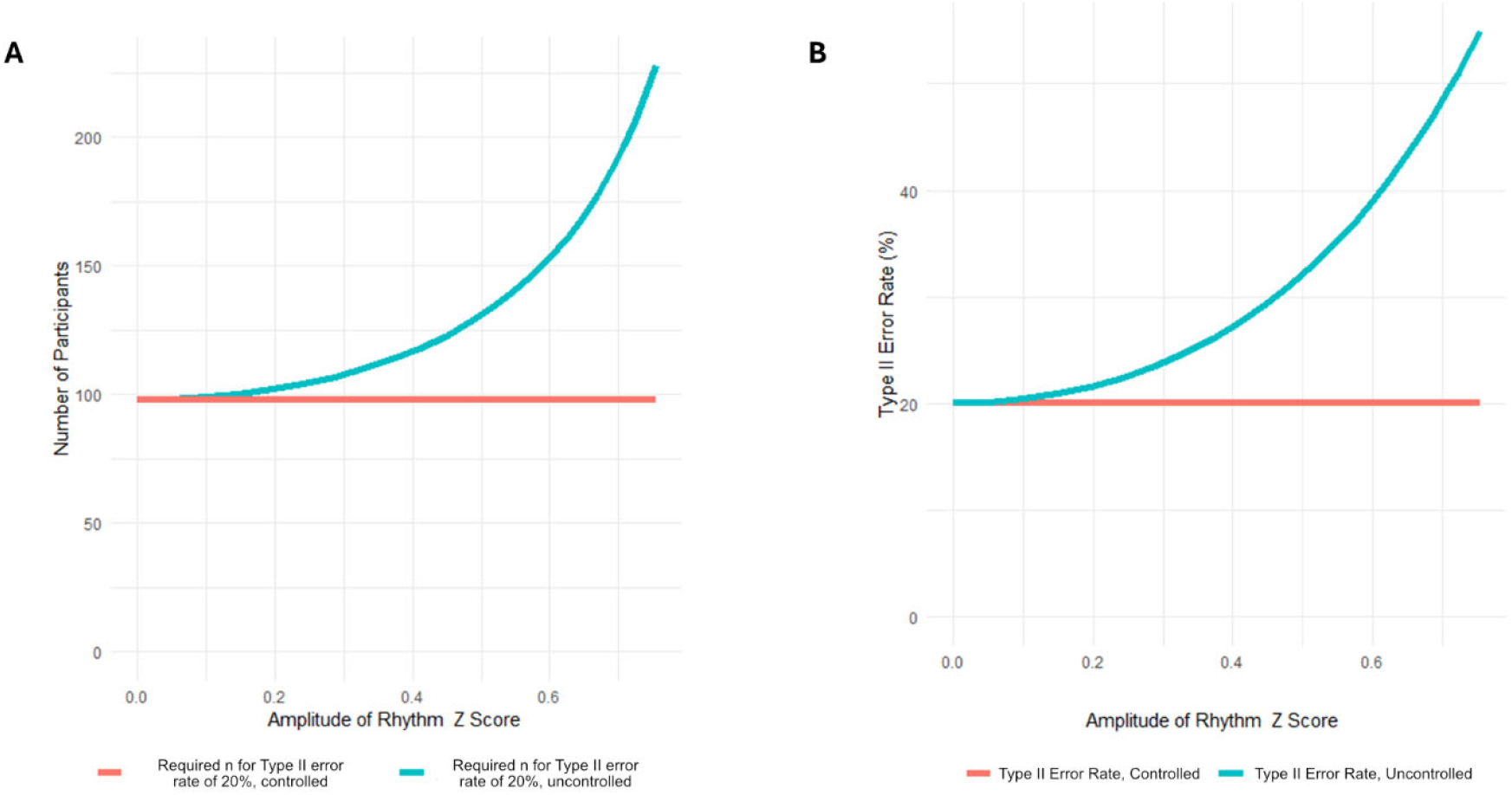
(A) Relationship between required *n* and amplitude of rhythm to achieve a 20% Type II error rate (B) Type II error rate with fixed *n* (number of participants) and an increasing independent cosinor rhythmic component

## Discussion

Temporal variation is a key aspect of physiology. Daily variation in the transcriptome and metabolome of human tissues and fluids have been described, ^27–29^ but rhythms of human proteomic data are poorly understood. Here we provide an analysis of the influence of temporal variation in the human serum proteome. Overall, 13 out of the 73 proteins meeting QC thresholds were identified as rhythmic, 6 being statistically significant at 24 hours, and 9 being statistically significant at 12 hours (including 2 rhythmic at both). The proportion of proteins identified as rhythmic was 18%. This is broadly consistent with a recent larger proteomics study which found that 15% of analysed proteins exhibited significant daily rhythmicity, ^13^ and is also concordant with metabolomics studies suggesting a range of 15% to 20% of features exhibiting circadian rhythmicity in constant routine conditions (in non-constant / entrained conditions this can be higher). ^9^ Rhythmicity of many of the biological functions captured here is already well-described, with immunoglobulins having previously been described as having a circadian component. ^30–32^ As well as immunoglobulins, this work also highlights a number of protein clusters and pathways as rhythmic, such as the coagulation and the complement cascade as well as apolipoproteins. Several of the individual proteins here have previously been identified as rhythmic, in particular the apolipoproteins APOA2, APOC3 and APOE. ^13,33^ Within the complement and coagulation cascades, FIBA, FIBB and KNG1 have also previously been reported as rhythmic. ^33^ The other proteins - ZA2G, ITIH2, PLG and CFAH - are to our knowledge identified for the first time as rhythmic in this work.

Circadian variation of fibrinolytic and complement activity in blood is a well-described phenomenon, ^34^ and has for example been reported to contribute to increased risk of cardiovascular events in the morning. ^35^ The proteins identified here have also been closely associated with conditions known to dysregulate circadian rhythms such as Alzheimer’s disease. For example, CFH within the complement cascade has been identified as a biomarker of Alzheimer’s, ^36,37^, as have the fibrinolysis pathway proteins FGA and FGB, ^38,39^ and indeed as has AZGP1. ^40^ Three apolipoproteins were also identified as rhythmic; the genes governing the expression of many of this family of proteins have previously been demonstrated to show rhythmic expression ^41^ and these results are also consistent with lipid metabolism more broadly having a circadian component; ^42 40^ Apolipoproteins have also been directly linked to Alzheimer’s disease; especially with regard to APOE4. ^43,44^ It should be noted that in some cases, where rhythmic proteins are identified as biomarkers of a condition, there is a risk that they are in fact biomarkers of generic circadian dysregulation, and are not specific to the disease in question. This is a known issue in multivariable analyses such as proteomics or metabolomics, especially when machine learning algorithms are trained on idealised case-control datasets of disease versus healthy participants. ^45^

In terms of limitations, the work presented here is a small case study reviewing only a limited number of proteins. It should be noted that recent analyses of proteomic rhythmicity have identified a much wider range of proteins with a circadian component. ^13,33^ Nonetheless, the statistical impact on biomarker or pathway identification shown in this case study is relevant for any rhythmic protein, transcript, metabolite, or other chronobiological biomarker of interest. Therefore, we can view chronobiology as representing a potential ‘omics confounder. As presented here, controlling for rhythmic variation allows for a reduction of sample size *n* whilst maintaining statistical power, whilst conversely failing to control for rhythmicity *ceterus paribus* increases Type II error rates. Therefore, rhythmicity represents both a meaningful opportunity (for improved statistical power at relatively low cost) in biomarker research, as well as a cost (in the form of increased error rates and reduced reproducibility), when not controlled for.

We see the following suggested measures as helpful in aiding the reader’s understanding of biomarker research in the context of chronobiology, reducing the risks of Type I and Type II errors, and improving reproducibility.

1. The ideal is to control for rhythmicity as part of the study design, especially when the biomarkers being investigated have a known rhythmic component or in untargeted studies that may have a rhythmic component. At the simplest level, time of day of sampling could be controlled; a better approach is to adjust for circadian phase with the gold-standard being the alignment to DLMO, albeit this will be beyond the scope of the vast majority of projects. Alternatively, single point-in-time methods could be used to calculate and adjust to a DLMO time. ^46^
2. If rhythmicity cannot be controlled for during sampling, statistical power calculations during the study design phase should acknowledge the impact of rhythmicity, and the necessary increase in *n* or the reduction in β.
3. We also recommend that time of day of sampling be considered as part of a study’s metadata, and should be reported, alongside for example, the p-value for any difference between case and controls’ time of sampling. This would also be of benefit in biobank data, as researchers would have the option of selecting time-of-day matched data to control for rhythmicity.
4. Finally, where rhythmicity in any identified biomarkers has previously been reported, this should be noted, especially in studies reviewing biomarkers, pathways, or conditions which are known to be influenced by the human timing system.

In conclusion, the rhythmic protein expressions shown in this work demonstrate the potential for confoundment; biological rhythmicity therefore presents a study design problem for case-control experiments in proteomics and other ‘omics platforms. Whilst aligning rhythms to an individual’s circadian phase using DLMO to remove the confounder is likely to be too costly for the majority of studies, adoption of the recommendations suggested in this work would mitigate against the risks of Type I and Type II errors and improve reproducibility of biomarker identification.

## Author contributions

D.R.v.d.V., D.J.S., and J.D.J. designed the clinical study. C.M.I. designed the diets. C.M.I. and H.H. managed sampling and clinical data collection. L.G. managed the proteomic data acquisition. M.S. analyzed the proteomics data and wrote the manuscript. All authors reviewed the manuscript.

## Data and code availability

The raw mass spectrometry data reported here will be available at the PRIDE repository. All code used in the preparation of this manuscript used open access libraries without modification.

## Acknowledgments

The authors thank staff at the Surrey Clinical Research Facility for their help with the clinical study and Ms. Jenny Spinks, chronobiology technician, for conducting the melatonin assays. The authors also wish to thank Dr. Thomas PM Hancox and Dr. Anthony Onoja for their advice on the theoretical and statistical underpinnings of the manuscript, as well as Waters Corporation for their support with the proteomics resourcing, methodology and analysis.

## Supplementary Material

**Table S1:**
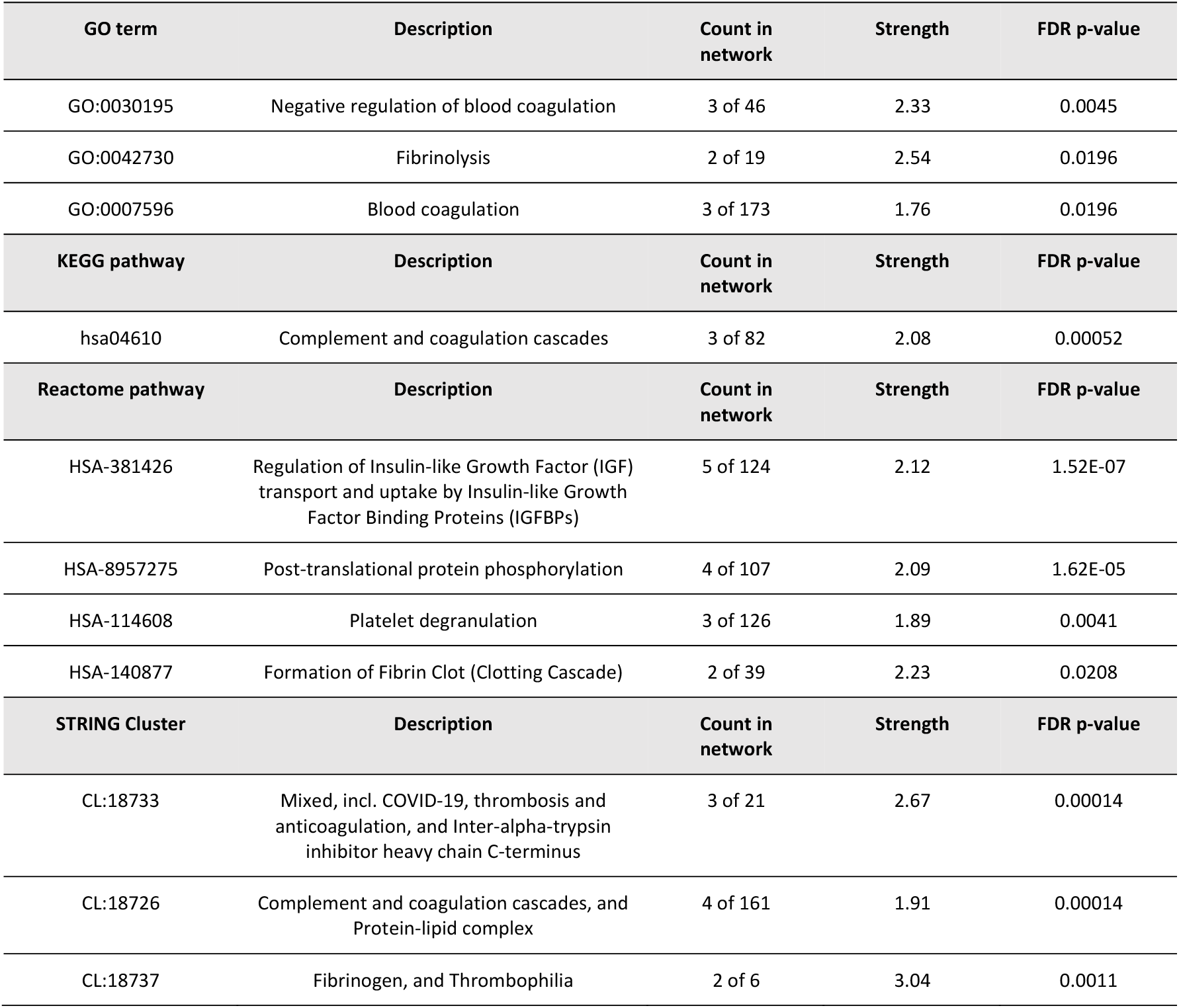
Proteins (genes) with 12 hour rhythm analysed by STRING, pathway outputs.

| GO term | Description | Count in network | Strength | FDR p-value |
| --- | --- | --- | --- | --- |
| GO:0030195 | Negative regulation of blood coagulation | 3 of 46 | 2.33 | 0.0045 |
| GO:0042730 | Fibrinolysis | 2 of 19 | 2.54 | 0.0196 |
| GO:0007596 | Blood coagulation | 3 of 173 | 1.76 | 0.0196 |
| KEGG pathway | Description | Count in network | Strength | FDR p-value |
| hsa04610 | Complement and coagulation cascades | 3 of 82 | 2.08 | 0.00052 |
| Reactome pathway | Description | Count in network | Strength | FDR p-value |
| HSA-381426 | Regulation of Insulin-like Growth Factor (IGF) transport and uptake by Insulin-like Growth Factor Binding Proteins (IGFBPs) | 5 of 124 | 2.12 | 1.52E-07 |
| HSA-8957275 | Post-translational protein phosphorylation | 4 of 107 | 2.09 | 1.62E-05 |
| HSA-114608 | Platelet degranulation | 3 of 126 | 1.89 | 0.0041 |
| HSA-140877 | Formation of Fibrin Clot (Clotting Cascade) | 2 of 39 | 2.23 | 0.0208 |
| STRING Cluster | Description | Count in network | Strength | FDR p-value |
| CL:18733 | Mixed, incl. COVID-19, thrombosis and anticoagulation, and Inter-alpha-trypsin inhibitor heavy chain C-terminus | 3 of 21 | 2.67 | 0.00014 |
| CL:18726 | Complement and coagulation cascades, and Protein-lipid complex | 4 of 161 | 1.91 | 0.00014 |
| CL:18737 | Fibrinogen, and Thrombophilia | 2 of 6 | 3.04 | 0.0011 |

**Table S2:** Proteins (genes) with 24 hour rhythm analysed by STRING, pathway outputs.

| GO term | Description | Count in network | Strength | FDR p-value |
| --- | --- | --- | --- | --- |
| GO:0034382 | Chylomicron remnant clearance | 2 of 5 | 3.12 | 0.0021 |
| GO:0043152 | Induction of bacterial agglutination | 2 of 6 | 3.04 | 0.0021 |
| GO:0032489 | Regulation of Cdc42 protein signal transduction | 2 of 8 | 2.91 | 0.0029 |
| GO:0090209 | Negative regulation of triglyceride metabolic process | 2 of 10 | 2.82 | 0.004 |
| GO:0031639 | Plasminogen activation | 2 of 11 | 2.78 | 0.0045 |
| KEGG pathway | Description | Count in network | Strength | FDR p-value |
| hsa04610 | Complement and coagulation cascades | 4 of 82 | 2.2 | 1.69E-06 |
| hsa04979 | Cholesterol metabolism | 2 of 48 | 2.14 | 0.0158 |
| hsa05150 | Staphylococcus aureus infection | 2 of 86 | 1.88 | 0.0327 |
| hsa04611 | Platelet activation | 2 of 122 | 1.73 | 0.0487 |
| Reactome pathway | Description | Count in network | Strength | FDR p-value |
| HSA-8964058 | HDL remodeling | 2 of 10 | 2.82 | 0.0116 |
| HSA-8963901 | Chylomicron remodeling | 2 of 10 | 2.82 | 0.0116 |
| HSA-8963888 | Chylomicron assembly | 2 of 10 | 2.82 | 0.0116 |
| HSA-372708 | p130Cas linkage to MAPK signaling for integrins | 2 of 15 | 2.64 | 0.0116 |
| HSA-354194 | GRB2:SOS provides linkage to MAPK signaling for Integrins | 2 of 15 | 2.64 | 0.0116 |
| STRING Cluster | Description | Count in network | Strength | FDR p-value |
| CL:18726 | Complement and coagulation cascades, and Protein-lipid complex | 6 of 161 | 2.09 | 1.55E-09 |
| CL:18737 | Fibrinogen, and Thrombophilia | 3 of 6 | 3.22 | 1.20E-06 |
| CL:18728 | Complement and coagulation cascades, and Positive regulation of opsonization | 4 of 109 | 2.08 | 9.96E-06 |
| CL:18956 | Lipoprotein particle, and Assembly of active LPL and LIPC lipase complexes | 2 of 46 | 2.15 | 0.0305 |

